# Eye tracking-based estimation and compensation of chromatic offsets for multi-wavelength retinal microstimulation with foveal cone precision

**DOI:** 10.1101/599464

**Authors:** Niklas Domdei, Michael Linden, Jenny L. Reiniger, Frank G. Holz, Wolf M. Harmening

**Affiliations:** Department of Ophthalmology, University of Bonn, Germany

## Abstract

Multi-wavelength ophthalmic imaging and stimulation of photoreceptor cells requires consideration of chromatic dispersion of the eye, manifesting in longitudinal and transverse chromatic aberrations. Current image-based techniques to measure and correct transverse chromatic aberration (TCA) and the resulting transverse chromatic offset (TCO) in an adaptive optics retinal imaging system are precise, but lack compensation of small but significant shifts in eye position occurring during in vivo testing. Here we present a method that requires only a single measurement of TCO during controlled movements of the eye to map retinal chromatic image shifts to the image space of a pupil camera. After such calibration, TCO can be compensated by continuously monitoring eye position during experimentation and by interpolating correction vectors from a linear fit to the calibration data. The average change rate of TCO per head shift and the correlation between Kappa and the individual foveal TCA are close to the expectations based on a chromatic eye model. Our solution enables continuous correction of TCO with high spatial precision and avoids high light intensities required for re-measuring TCO after eye position changes, which is necessary for foveal cone-targeted psychophysical experimentation.

## Introduction

Adaptive optics scanning laser ophthalmoscopy (AOSLO) coupled with microstimulation techniques enables imaging and simultaneous functional testing of targeted human photoreceptors in vivo. This approach was recently employed to study single cone photoreceptor function [1,2], retinal circuitry [3,4], color vision [5,6], and sensitivity changes during retinal disease [7,8].

The specific techniques employed in these studies, using two (or more) beams of light of different wavelengths for imaging and stimulation, are faced with a particular challenge arising from the chromatic dispersion characteristics of the human eye. Due to dispersion, light of different wavelengths will be focused axially displaced (termed longitudinal chromatic aberration, LCA) [9], and if incident at an oblique angle, focus points will also be laterally displaced (termed transverse chromatic aberration, TCA) [10].

The magnitude of LCA is relatively consistent across the population [9,11] and across the field of view [12,13], thus it can be corrected sufficiently for most eyes by adjusting the relative vergence between beams of different wavelength. In an AOSLO system, this is currently achieved by a coaxial displacement of the light sources’ entry points.

By setting the light sources at different distances, transverse chromatic offset (TCO) is induced in the eye. These lateral offsets in focus position on the retina have two origins. One is the imperfect axial positioning of the two (or more) beams. The second is of ocular origin, and closely related to TCA. When the eye is moved laterally in front of the displaced beams, their focus location will move laterally on the retina as a linear function of eye position, akin to a chromatic parallax. The magnitude of this offset is identical to TCA but opposite in sign [14].

In an AOSLO, TCO can be directly measured by spatially registering simultaneously recorded retinal images with the two (or more) wavelengths. This method allows a determination of TCO of the order of arcseconds but requires high light intensities massively bleaching the photopigment and leading to strong fluctuations in retinal adaptation [14]. Therefore, continuous or repeated measurements of TCO during psychophysical sessions are unfeasible.

In earlier studies, TCO was thus statically compensated by assuming no lateral eye motion between measurements before and after an experimental run. This was only valid as long as the cone photoreceptor diameter at the targeted eccentricities was large enough to allow typical residual head and eye movement which would displace the stimulus within the cone’s diameter [1,3,5,6]. For stimulation of the smallest cones at the foveal center or rods, however, this approach is no longer sufficient.

A chromatic eye model by Thibos et al. [15–17] predicts a simple linear correlation between TCO and eye position offsets in front of the AOSLO beams, depending on the wavelengths, described by:

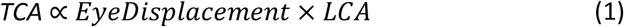

This relationship was experimentally confirmed in previous studies [14,18], and applied to a method to infer TCO from pupil position, yet without focusing on the precision necessary to target single cones [19]. We here employ high-resolution eye tracking to demonstrate that transverse chromatic offsets can be compensated in real time to ensure cell sized precision during single cell psychophysical experiments on cones of the central fovea or rods.

## Materials and Methods

To map lateral eye position changes to transverse chromatic offset changes, we employed an adaptive optics scanning laser ophthalmoscope (AOSLO) to measure image-based retinal TCO [14] during controlled movements of the head and simultaneous eye tracking. Following such calibration, eye tracking-based TCO estimates were validated in a psychophysical experiment. A detailed description of our AOSLO system is given in [20]. TCO calibrations were performed in fourteen participants (9 female, 5 male) with no known vision abnormalities. Three participants (1 female, 2 male) took part in the subjective experiments. Mydriasis and cycloplegia were induced by instilling one drop of 1%Tropicamide 15 min before the beginning of a session. For each participant, a holder fixed to a custom dental impression (bite bar) was used to immobilize and control the position of the head during imaging. Written informed consent was obtained from each participant and all experimental procedures adhered to the tenets of the Declaration of Helsinki, in accordance with the guidelines of the independent ethics committee of the medical faculty at the Rheinische Friedrich-Wilhelms-Universität of Bonn.

### 2.1 Eye tracking camera

In order to accurately track the position of a subject’s eye in front of the AOSLO beam, we mounted a video camera coaxially to the AOSLO imaging and stimulation beams by means of a cold mirror (DMLP900L, Thorlabs) (Fig.1). The camera was a 752 × 480 pixel CMOS sensor (DMK 23UV024, The Imaging Source) with a 50 mm, f/1.26 objective lens (TCL 5026 5MP, The Imaging Source). A central 640 × 480 pixels subfield of the sensor was used in our custom eye tracking software. Camera focus was set to a fixed working distance of 350 mm. By judging the sharpness of the pupil image within the eye tracking user interface, eyes could be positioned at an approximate distance coinciding with a conjugate pupil plane of the AOSLO beam. Image magnification of the eye tracker was 0.030 mm per image pixel in the pupil plane. As a result of back-scattered light from the imaging beam (7.2 mm diameter), the camera captured an image of the retro-illuminated pupil on an otherwise dark background (Fig. 1C, D). The imaging beam also produced a bright visible reflection on the cornea front surface, the first Purkinje Image, referred to as Purkinje image in the following. The bright image pixels of the Purkinje image were used to track the eye’s lateral location. Because of the coaxiality between camera and illumination, we used the horizontal offset between the pupil’s center and the Purkinje image to calculate the horizontal component of angle Kappa, κ, (the angular subtense between visual and pupillary axis) as follows:

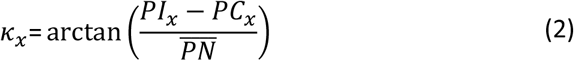

**Fig. 1.**
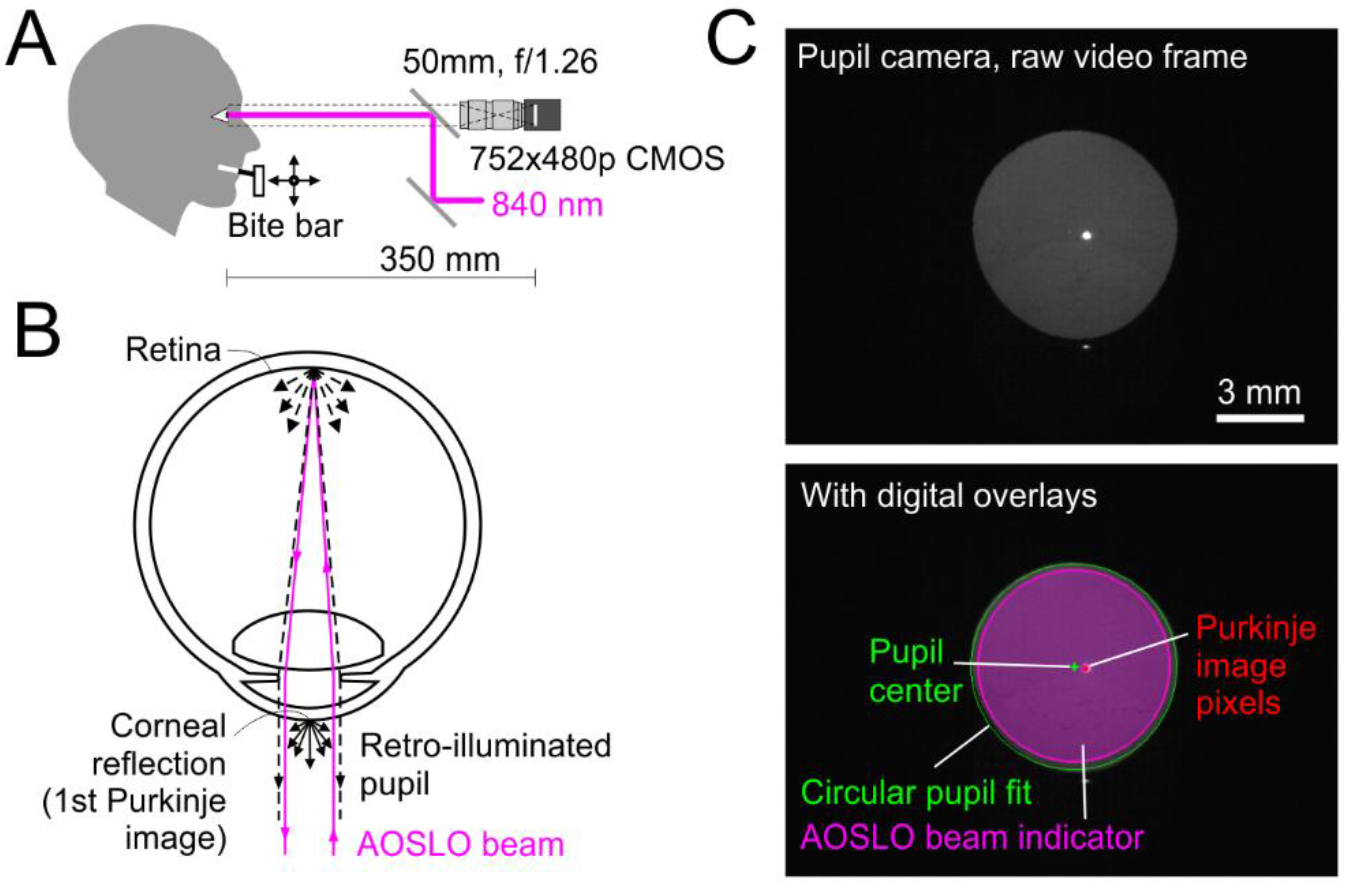
On-axis eye tracker with AOSLO as light source. A: Side-view, to scale. The subject’s head could be moved via XYZ-microdrives attached to a bite bar. B: Using the 840 nm beam of the AOSLO as light source, retinally back-scattered light illuminates the pupil from within the eyeball. The 1st Purkinje image was used to track the eye’s position relative to the AOSLO beam. “N” marks the first Nodal point of the eye and defines the intersection between pupillary and visual axes. Note that the camera is aligned with the visual axis. C: Pupil and Purkinje image could be tracked with high precision (1 image pixel equaled 30 μm in the pupil plane). During operation, digital overlays could be displayed to aid positioning during calibration.

PIx defined as the horizontal Purkinje image coordinate, PCx as the horizontal pupil center coordinate, and 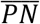 as the distance from the entrance pupil to the Nodalpoint, which was assumed to be 4 mm [21]. The vertical component of Kappa was calculated in the same way. Additionally, we calculated the Hirschberg ratio (the inverse of the displacement of the Purkinje image from pupil center per degree of eye rotation) in four participants during controlled gaze shifts. The average Hirschberg ratio across four participants was 13.3 ± 0.7 °/mm, which corresponds to an average 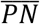 of 4.2 mm. The individual values of 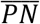 were used to calculate the participant’s Kappa more precisely and estimate TCO based on LCA and Kappa.

### 2.2 Eye tracking software

A custom C++ program based on previously described algorithms [22,23] was written to display, track, and measure certain image features visualized by the eye tracking camera (Fig. 1D). Each of the following metrics was computed for every new frame with a refresh rate of 29.87 Hz. The pupil was fitted by a circle to determine its center and diameter. The Purkinje image coordinate was specified by the center of a second circle fit to the brightest pixels of the corneal reflection. Pixel detection thresholds for the circle fits were adjusted manually at the beginning of each session to ensure the best fits at the current image quality. To assist eye alignment, an AOSLO beam indicator marking the perimeter of the imaging beam was added. AOSLO beam position was captured at the beginning of every session based on the camera image of a white paper illuminated by the imaging beam in the pupil position. While running a TCO calibration sequence, all data was written to a log file containing frame number, time stamp, pupil center coordinates, Purkinje center coordinates, and pupil diameter.

### 2.3 TCO measurement

A detailed description of the method to objectively measure TCO with an AOSLO is given in [14]. In short, a video is recorded containing interleaved information of both imaging and stimulation light channels (here, 840 nm and 543 nm) in a subfield of the imaging raster (192 × 128 pixel). Images recorded in both channels were spatially cross-correlated frame by frame to compute the positional shift between the imaging light and stimulation light on the retina. Because this method is image-based, it requires video footage with resolved retinal structure which can be a limiting factor at the central fovea. As a side note, we observed that image structure moved if the confocal pinhole in front of the light detector is moved. As a consequence, TCO readings will change, although no changes occurred on the retina. To control for this additional system-related chromatic offset, we centered the confocal pinhole position on the beam’s point spread function by optimizing image brightness and contrast ahead of each session.

### 2.4 Calibration procedure

To correlate image-based measurements of TCO to eye position changes, a 20 sec dynamic calibration sequence was performed. During recording, the head (and therefore the eye) was moved in a somewhat random pattern with an extent of about ± 0.2 mm in each direction by manually turning the knobs of a XY micrometer stage attached to the bitebar. Meanwhile, the participant fixated on a target ensuring video acquisition of the same retinal location which was at ~0.4 degree eccentricity. Reminiscent of a clapperboard in film industry, we modulated the AOSLO beam through three quick full on and off cycles to flash the imaging beam at the beginning and at the end of the recording, a signal which could be accurately assigned to a single frame in both data streams. TCO video data and eye tracking data were processed offline with a custom Matlab (Mathworks, Inc.) software in three steps (Fig. 2):

1. Synchronization and interpolation: the two independent data streams were synchronized based on the flash-sequence flagging “start” and “end” by downsampling TCO data via linear interpolation - due to slightly differing framerates (TCO: 30.00 Hz, Eye tracker: 29.87 Hz) - to assign an individual TCO data sample to each Purkinje image position (Fig. 2A).
2. Image noise removal: changes between consecutive TCO samples larger than 2 pixels (threshold determined empirically) were regarded as image noise and removed from further analysis (Fig. 2B). Single samples with change rates below threshold that lay in immediate neighboring blocks of samples flagged as noise were also removed.
3. TCO-tracking correlation: the linear regression of TCO and eye tracking data for both horizontal and vertical component was computed and then used to estimate TCO from eye position. The linear regression (for horizontal eye shifts) was in the form of:

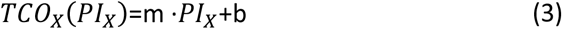

wherePIx is the Purkinje’s image X-coordinate, m and b are the slope and a constant offset of the linear regression and TCOx is the resulting retinal image offset (Fig. 2C). The estimate for the vertical component was computed accordingly.

**Fig. 2.**
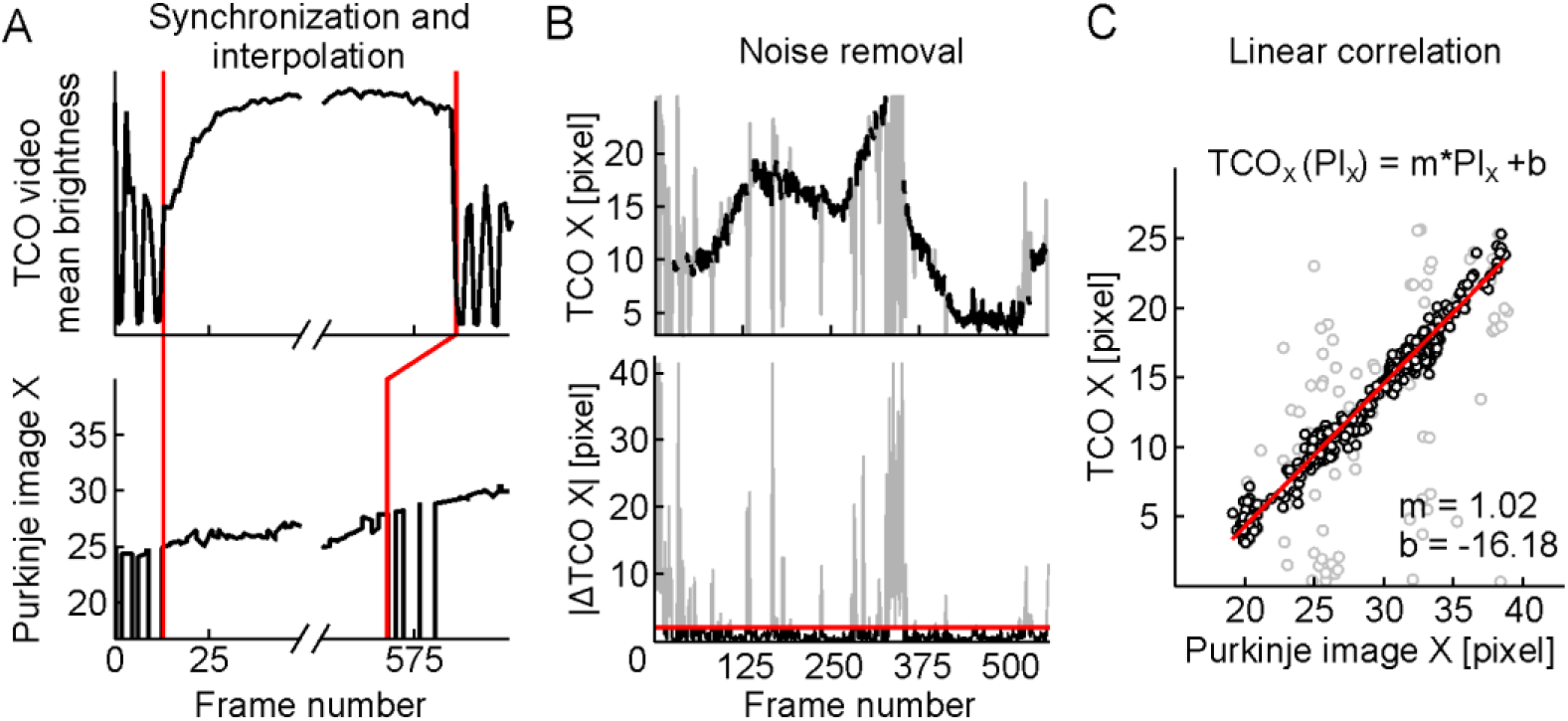
Calibration processing steps. TCO and eye tracking data were recorded simultaneously while the operator moves the participant’s eye in front of the system. A: For synchronization, three quick full on and off cycles flashed the imaging beam (red lines). Due to slight differences in sampling rate, TCO data was down-sampled via linear interpolation. B: Larger TCO data excursions (grey) were removed by thresholding for frame by frame TCO sample changes, Δ TCO (bottom graph, red line marks 2-pixel-threshold). C: Computation of the correlation between TCO and Purkinje image position by least-squares linear fit. The resulting function (shown as inset) was used to continuously estimate TCO based on eye tracking data.

### 2.5 Experimental validation

For an objective validation of our eye tracking-based TCO estimation, we consecutively recorded 20 calibration sequences in one eye. We then used the linear regression function found in the first sequence and compared the measured TCO data at each eye position with the estimated TCO data by plugging in the eye position data in the calibration function for all following sequences. This was repeated for all 20 runs. Precision was estimated by using the x and y component of each frame’s error as a coordinate in a two-dimensional displacement histogram. Repeatability of the procedure was measured by comparing the TCO coordinates across all 20 calibrations by solving equation 3 for a single (average) eye position.

We validated TCO estimation in a psychophysical experiment with three participants. The participant’s task was to manually align their head in front of the AOSLO beam such that the features of a two-color centroiding stimulus produced in the AOSLO imaging raster were aligned. The stimulus feature offsets (a 2.3 arcmin dot of 543 nm light shown within a black target within the 840 nm imaging raster) were randomly chosen from sixteen offsets in a 4-by-4 grid with 0.4 arcmin spacing. Each offset was presented 6 times, for a total of 96 trials of subjective alignments. By the push of a button, the participant confirmed the correct alignment, the current Purkinje image location was stored, and a new stimulus offset presented. The predicted eye position based on the TCO calibration sequence and chromatic stimulus offsets were compared with the actual eye position at each trial.

## Results

We performed TCO eye tracking calibrations in a total of 14 eyes, with multiple (3-20x) repetitions in all eyes, producing a total of 62 calibration sequences. We found that in order to collect sufficient data points (of both, image-based TCO and eye tracking data) during the 20 sec calibration, the most efficient head movement pattern resembled a square or circle. Figure 3 shows four example calibration sequences from three eyes with the produced raw data and subsequent regression analyses. To review calibration success, three different metrics were displayed after each calibration sequence, which also guided repeat decisions if a sequence had failed (e.g. due to insufficient image quality):

1. Percentage of clean data samples,
2. Eye movement extent in both horizontal and vertical direction given as distance of the center 80%quantiles of all clean location samples (Q80),
3. Coefficient of determination of the linear regression to the data (R2).

**Fig. 3.**
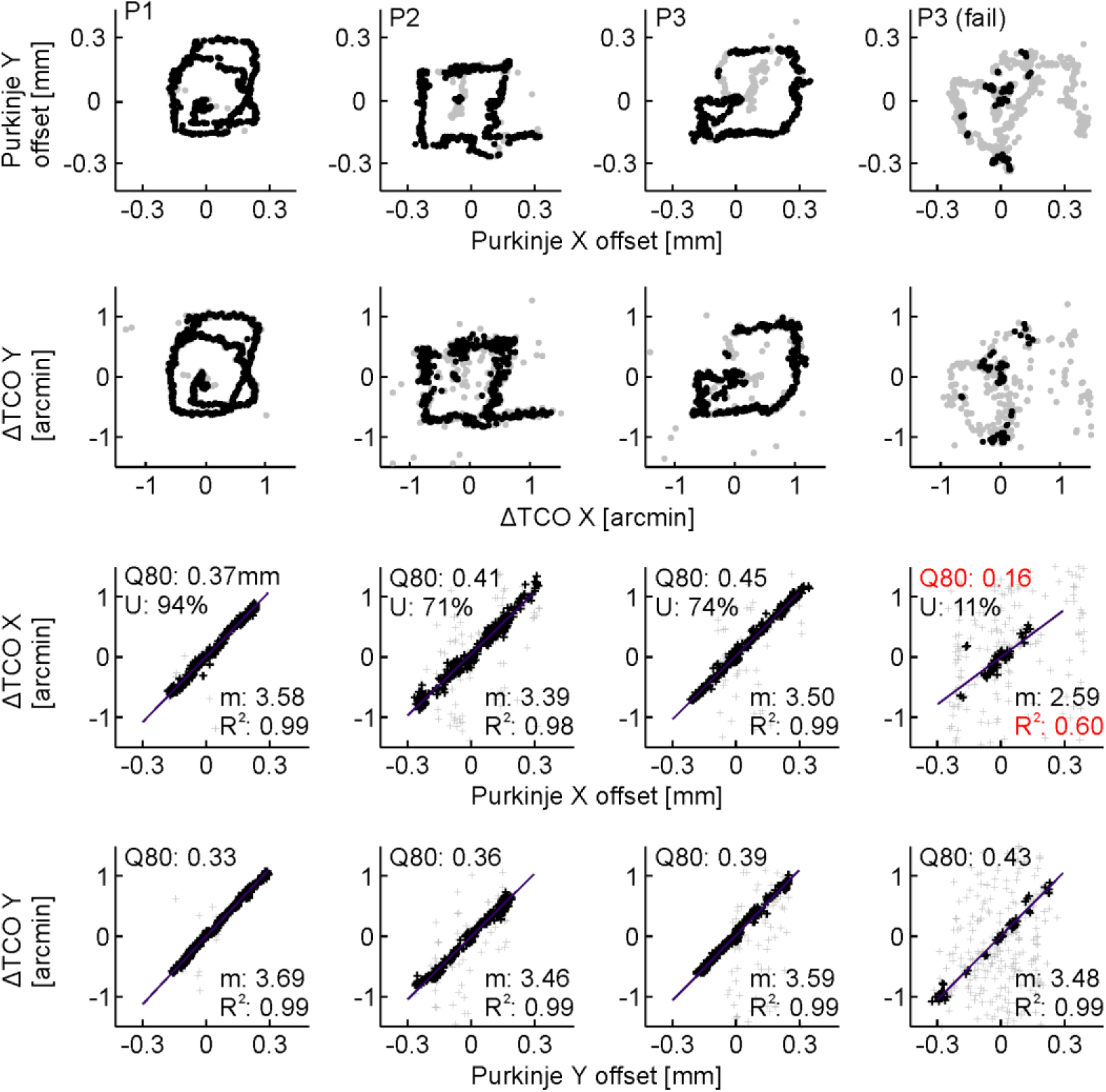
Calibration sequences recorded in three participants (P1, P2, P3). Top row: Captured Purkinje image positions during calibration. 2nd row: TCO data of the same calibration sequence, based on video data recorded at 0.4 degree eccentricity. 3rd and 4th row: Linear regression for horizontal and vertical eye shifts, respectively. Percent samples used (U), range of data points (Q80) and goodness of fit (R2) are metrics chosen to support the operator’s decision whether a calibration has to be repeated (an example is shown in the 4th column). Black data points are the usable samples after interpolation and removing data noise, grey data points represent the data samples flagged as noise (see Methods for details).

When image quality decreased, the TCO computation algorithm produced an increasing number of noisy data (Fig.3, 4th column). We found that in order to ensure a reliable calibration, the percentage of used samples should exceed 33%. The second metric, Q80, should exceed 0.3 mm in vertical and horizontal direction for a successful calibration (see an example of this metric taking effect in Fig. 3, last column). The third metric, R2, was mostly greater than 0.96 and calibration sequences with R2 > 0.9 were considered to be sufficiently accurate.

Based on a chromatic model eye describing the relationship between LCA, TCA and eye position [17], we expected the correlation between eye lateral position and TCO to be linear (equation (2)). We found the slope, m, of the linear regression was constant across eyes but different for horizontal and vertical shifts. Considering the median slope of three consecutively recorded calibrations in each eye, the mean slope for horizontal eye shifts was 3.55 ± 0.08 arcmin/mm and 3.43 ± 0.12 arcmin/mm for vertical eye shifts across subjects (Fig.4 A). By pairing horizontal and vertical data for each eye, this difference was significant (p = 0.02, Wilcoxon signed rank test). The absolute offset, b, of the linear regression was found to vary across participants, reflecting an idiosyncratic component of TCA (Fig. 4B). Because we also tracked the pupil’s center during calibration we could compute TCO values when the pupil was computationally centered on the beam (Fig. 4C).

**Fig. 4.**
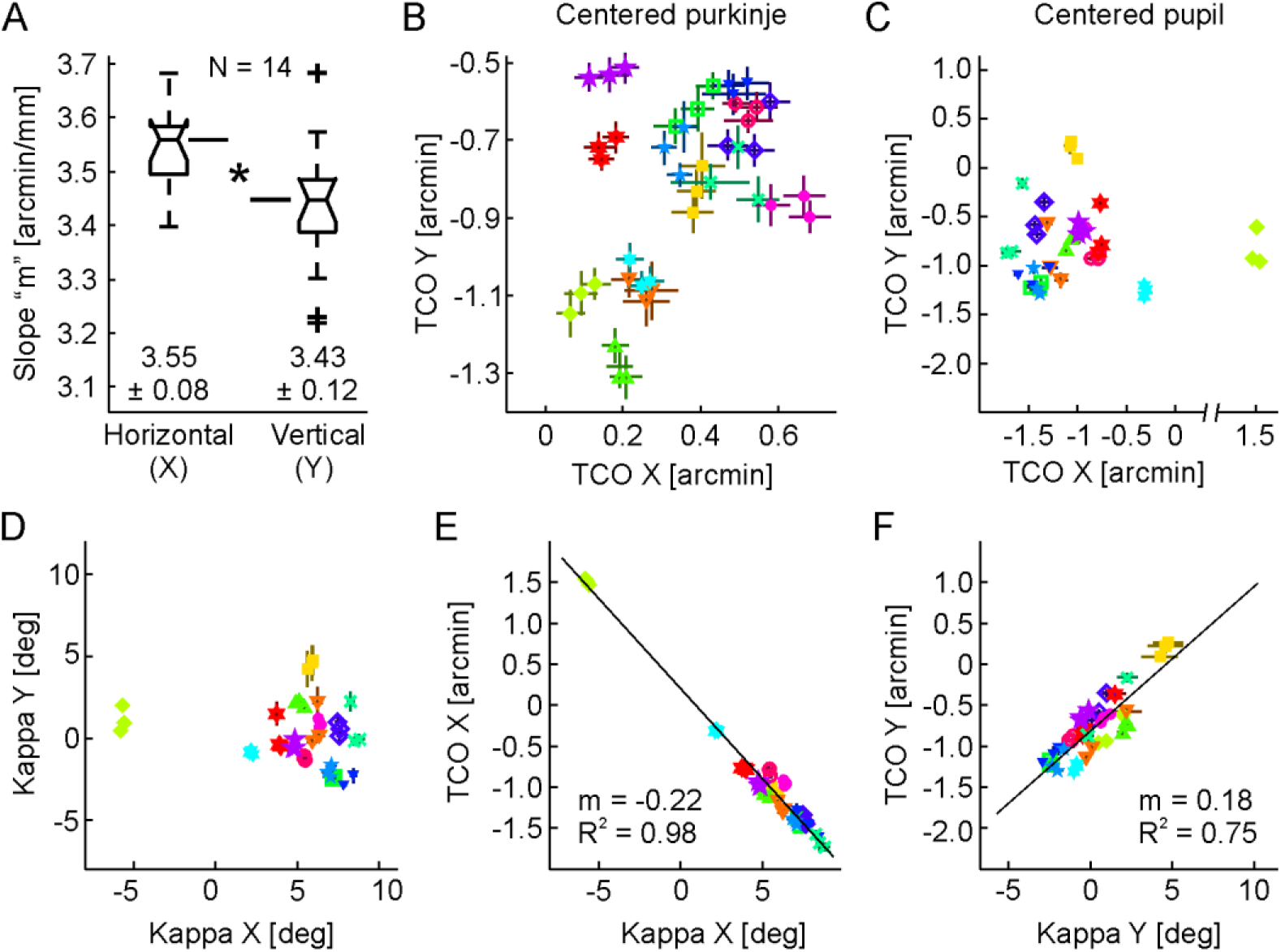
TCO data across 14 eyes of 14 participants. A: Boxplot of the median calibration slope for each eye (3 repeats each). The average horizontal correlation slope across eyes was 3.55 ± 0.08 arcmin/mm, the average vertical correlation slope 3.43 ± 0.12 arcmin/mm. This difference between horizontal and vertical slope was significant (p = 0.02, Wilcoxon signed rank test). B: Calculated TCO for each run with a Purkinje position centered on the AOSLO beam. C: Calculated TCO for a centered pupil position. The one left eye included in this data set shows an absolute TCO with in versed sign (green diamonds). D: Angle Kappa for all eyes. E: Correlation of horizontal Kappa and horizontal TCO of the centered pupil. F: Correlation of vertical Kappa and vertical TCO of the centered pupil. For plots B-F, vertical and horizontal bars mark the 0.5 quantile of the calibration function (TCO) and the standard deviation (Kappa). Different symbols mark different eyes. If no error bar is visible, the error is smaller than the symbol.

For a centered Purkinje position, individual TCOs were positioned close to each other, with an extent of 0.8 arcmin from lowest to highest value for both dimensions. The range of individual TCO values broadened when calculated for a centered pupil: TCO values for the right eyes spread within a range of 1.5 arcmin. TCO for the one left eye tested had the opposite sign, as expected. Finally, we calculated angle Kappa for each participant (Fig. 4D) and found a strong correlation between the TCO calculated for the centered pupil position and Kappa (Fig. 4E,F). The linear regression of horizontal TCO with Kappa was defined by a slope of m = − 0.22 arcmin/deg (R^2^ = 0.98) and m = 0.18 arcmin/deg (R^2^ = 0.75) for the vertical dimension.

To estimate measurement error of our eye tracking based method, we compared 20 consecutively recorded calibration sequences of one eye with each other (Fig. 5). The 0.9 quantile of the framewise calculated difference between estimated and measured TCO was 1.04 pixel (0.10 arcmin) and 0.66 (0.07 arcmin) for horizontal and vertical axis, respectively (Fig. 5A,B). Exceptions were single events - presumably due to eye blinks or bad image quality - which could be as high as 2 pixel (0.2 arcmin) (Fig. 5B). By comparing each data set with each other in all possible pairwise combinations disallowing redundancy, 190 error sets containing a total of 88,266 error frames could be analyzed (Fig. 5C). As an estimate of precision we calculated the area containing 95 % of all data points in a plot of positional error in both spatial dimensions between estimated and measured TCO. The extent of this area was ± 0.15 arcmin along the horizontal and ± 0.12 arcmin along the vertical axis. The area containing 50 % of all positional errors had a width of ± 0.07 arcmin and a height of ± 0.05 arcmin. To analyze the repeatability of measurement, we first determined the average eye position across all 20 calibrations. This value was plugged in for each calibration function to calculate the TCO estimate for this position (Fig. 5D). The standard deviation for these 20 TCO estimates was 0.022 arcmin on the horizontal and 0.024 arcmin on the vertical axis. Across the three repeated calibrations in 14 participants we observed a median standard deviation for absolute TCO estimates with a centered Purkinje image of 0.029 arcmin (min=0.012; max=0.062) on the horizontal and 0.038 arcmin (min=0.013; max=0.070) on the vertical axis (Fig. 4B).

**Fig. 5.**
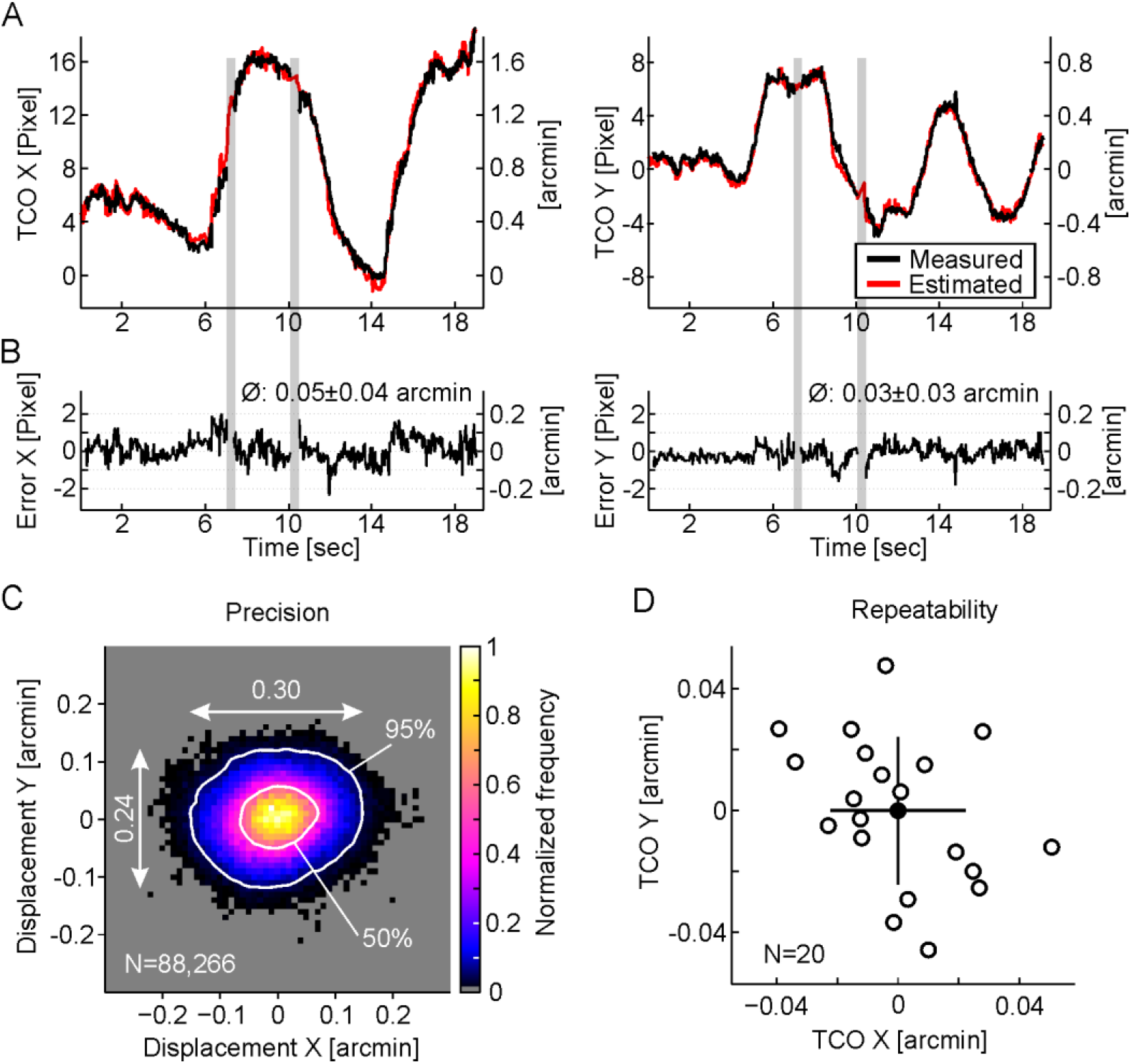
Error estimation. A: Exemplary comparison between measured TCO (black) and estimated TCO (red) following a single calibration sequence. Grey areas mark examples where image quality did not allow TCO measurement. B: Error of TCO estimation over time. Average (± std) estimation error was 0.05 ± 0.04 arcmin. Single events may exceed an estimation error of 1 pixel (1 pixel = 0.1 arcmin). C: To determine TCO estimation precision, 20 consecutive calibration sequences were validated against each other. The framewise displacement error was plotted in both spatial dimensions with one-tenth pixel resolution (1 block = 0.01 arcmin). 95 % of all displacement errors were within ±0.15 arcmin (horizontal) and ±0.12 arcmin (vertical). D: Repeatability was tested by using the average eye position in the 20 calibration functions (open circles). The standard deviation across all data is visualized via the extent of the horizontal and vertical bars (STDx = 0.022 arcmin/ STDy = 0.024 arcmin).

Finally, a subjective validation experiment addressed the psychophysical component of the TCO estimation. Three participants were asked to adjust their head position in front of the AOSLO beams to align a centroid stimulus containing controlled chromatic offsets (Fig. 6A). TCO estimation quality was assessed by comparing the applied chromatic stimulus offset with the resulting TCO estimated from the participant’s self-adjusted eye position (Fig. 6B). The average difference between calculated TCO from estimation and the actual applied offset was 2.49, 2.96, and 2.95 pixel (0.25, 0.30 and 0.30 arcmin) for P1, P2, and P3, respectively. Each participant showed an individual alignment strategy, resulting in clustering of certain displacement offsets. When pooling data across participants, no systematic direction for displacement errors was evident.

**Fig. 6.**
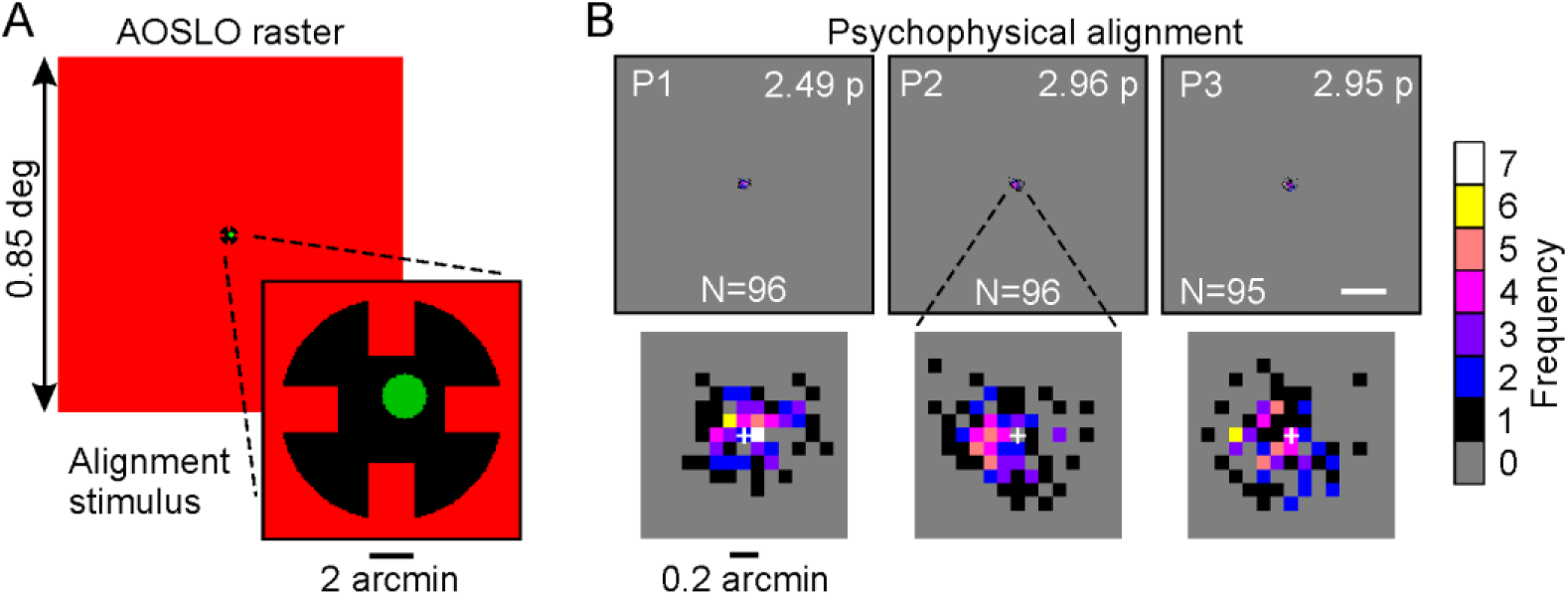
Subjective validation of our eye tracking based TCO estimation approach. A: AOSLO raster with stimulus to scale. The task was to center the green circle within the red notches. B: Results from all 3 participants in the same scale as the zoomed stimulus in A (scale bar = 2 arcmin). The mean difference between predicted position from estimation and subjective alignment was 2.49, 2.96, and 2.95 pixels (0.25, 0.30 and 0.30 arcmin) for P1, P2, and P3, respectively. There was no systematic direction for displacement errors. The enlargement below each panel in B displays the pixelwise TCO correction errors for each participant. The white cross marks the zero difference position.

## Discussion

Here, we present a video-based eye tracking approach to estimate transverse chromatic offsets (TCO) in an adaptive optics scanning laser ophthalmoscope (AOSLO) retinal microstimulator. The ultimate goal of this work was to compensate system- and eye-inherent chromatic offsets with a precision enabling targeting and stimulation of the smallest retinal photoreceptors, foveal cones and rods, without the need of direct and continuous TCO measurement, while at the same time allowing small head (and eye) movements to occur. By using a corneal reflection of the AOSLO beam as eye position beacon, we achieved TCO estimation with low noise, high repeatability and high spatial precision. We here discuss our principle of Purkinje image based positional tracking, the relationship between transverse chromatic aberrations (TCA), TCO, and the angle Kappa of the eye, and finally its usability and application for single photoreceptor stimulation.

Many eye trackers use pupil coordinates such as its center to report absolute eye position, and the pupil center has been shown to coarsely correlate with objective measures of TCA [14,18,19]. For precise estimates of TCA, however, the location of the TCA-inducing beams relative to the eye’s optics - and not its pupil - is more relevant [24]. So far, correlations of pupil position with chromatic offsets were not sufficiently accurate to allow for TCO compensation with photoreceptor resolution [19]. We argue that the pupil center is not an appropriate proxy of TCO, because the axis defined by the pupil center can shift with respect to the visual axis, e.g. during pupil constriction and dilation [25]. Typical pupil trackers determine the pupil’s center applying a circle fit, which is sensitive to errors because circularity of the pupil changes with pupil size. Micro fluctuations of pupil size are abundant, and the size changes dramatically during cycloplegia [26], a dynamic and reversible situation which state is usually uncontrolled for during microstimulation experimentation. In our approach, we use the first Purkinje image, the reflection of the front surface of the cornea, of the AOSLO beam as the tracking signal for eye translation changes. The location of this reflection depends on both the absolute position of the eye relative to the light source and camera as well as the rotational state of the eye. In relationship with the positional changes of the pupil center during eye rotation, it can therefore also be used as a gaze signal [27].

In our setup, with a static camera coaxially aligned to the light source, lateral head and eye shifts as well as rotation of the eye ball will move the Purkinje image. Purkinje image movement caused by gaze shifts could induce an error in our TCO estimation, because measures of TCO changes with retinal eccentricity [28]. Eye ball rotation occurs in two different cases and in both cases we used the Hirschberg ratio to estimate the magnitude of eye rotations induced errors. The individual Hirschberg ratios of four eyes measured with our eye tracker were 12.2 degree/mm, 13.5 degree/mm, 13.6 degree/mm, and 13.7 degree/mm (average 13.3 degree/mm (data not shown here)), in good accordance with values found in the literature [22,29]. In the first case, gaze changes occur due to the lateral head motion during calibration while the participant maintains visual fixation of a static visual target. Shifting the head by 0.4 mm in one direction will induce a rotation of the eyeball of 0.05 degree. This angle will move the Purkinje image about 0.0038 mm or 1/10 of a pixel in the camera image, and is thus negligible. In the second case, fixational eye movements during calibration or later compensation will shift the Purkinje image with a ratio of about 2.5 pixel per degree of gaze shift. Hence, to displace the Purkinje image by a full pixel the participant would have to rotate the eyeball by 0.4 degree. Typical fixational eye movements such as tremor and drift have an amplitude of about 0.01 degree, or 0.1 degree respectively [30]. Thus, only microsaccades could move the Purkinje image with detectable amplitudes. Because of the AOSLO’s inability to correct stimulus locations for the fast changes in retinal location during a microsaccade, however, TCO compensation would not be needed in that situation.

As hypothesized by a theoretical chromatic eye model, the relationship between ocular TCA and small displacements of eye position (< 2 mm) ought to be linear, and dependent on wavelength [15–17]. We found an average slope of 3.55 arcmin horizontal TCO change per mm eye shift, and 3.43 arcmin/mm for vertical TCO. This finding is in accordance with earlier reports (3 arcmin/mm, 3.5 arcmin/mm, and 3.86 arcmin/mm [14,18,19]), and matches the chromatic eye model, predicting a slope of 3.44 arcmin/mm for a LCA of 1.00 diopter when 840 nm and 543 nm light is used [15–17]. The residual variations we observed might be caused by individual differences in LCA. In our group of 14 participants, the observed difference between slopes of horizontal and vertical direction was significant, an observation which was also found in a similar study [19]. Following equation (1), such difference could be caused by differences in optical dispersion along the horizontal and vertical dimension of the eye. Although this kind of dispersion anisometry has not been reported before, it could be hypothesized that the dispersive characteristic of the eye also differs between horizontal and vertical meridians, because curvature and therefore refractive power of the cornea differs between those meridians [31,32].

We determined absolute TCO values in 14 participants with a centered Purkinje image (due to the fact that our light source, visual axes and camera are all coaxially aligned), and found values ranging within 0.6 arcmin along the horizontal and 0.8 arcmin along the vertical dimension. These ranges are clearly smaller than the ranges of absolute foveal TCA estimation reported by previous studies: about 1 arcmin [24,28], 2 arcmin [15,33], 4 arcmin [17] and 8 arcmin [34]. In all these studies, TCA was determined with a centered pupil. The range of absolute TCO for a centered pupil position in our study spanned 1.4 arcmin and 1.6 arcmin for horizontal and vertical dimension, including only right eyes. The closer range of absolute TCO - including the single left eye - for a centered Purkinje position can be explained by the fact that the foveal achromatic axis and the visual axis are closely related [21], and that our eye tracking camera was aligned with the visual axis. The remaining scatter could be caused by individual differences of the eye’s optics. The reason for our observation that the TCO values with a centered Purkinje do not vary around zero may be due to a minimal alignment difference between the imaging and stimulation light in the light delivery path, or - since the TCO measurement is image based - the IR and green detector positions.

To study the relationship between visual, pupillary and achromatic axes in more detail, we also measured individual Kappa angles in all eyes. The measured TCO values for a centered pupil where linearly correlated with Kappa (R^2^ = 0.98 for horizontal, R^2^ = 0.75 for vertical Kappa). This finding supports the chromatic eye model, where TCA is correlated with the angle Psi ψ (defined by pupil displacement and nodal point distance from the entrance pupil, like Kappa, here) [15,17]. This experimental confirmation has important implications for all ophthalmic applications where TCA is wished to be measured or corrected. First, we confirmed that the slope, m, of TCA per pupil displacement can be directly derived from individual LCA values of the eye. Second, if beams are centered in the pupil, a precise measurement of Kappa can be a direct measure for foveal TCA, once their relationship has been established in the system used, determining the offset b (equation (3)). We tested this in four participants, by recalculating Kappa for the individual Nodalpoint distance obtained from the participant’s Hirschberg ratio. In that situation, where the TCO estimation function based solely on Purkinje image and pupil center data, we would have made an average error of 0.13 ± 0.04 arcmin for horizontal TCO and 0.14 ± 0.07 arcmin for vertical TCO. Thus, only if a spatial precision on the order of single cones of the central fovea and rods is desired, a direct calibration as presented here would be required. Other applications like functional testing of pathologic areas of the retina in patients where direct TCO measure is unfeasible or unsuitable due to the high light intensities could utilize real time TCO compensation, simply by determining Kappa [7,8].

One of the important goals of the eye tracking based estimation of TCO was to increase estimation precision up to single foveal cone level. The cones of the central fovea are the smallest photoreceptor cells of the retina with an inner segment diameter of about 30 arcsec [35] comparable to the diameter of rods. With increasing eccentricity, cones increase in diameter with the result that positional errors caused by a participant’s residual head movements become more negligible (Fig. 7). Typically, participants show individual amounts of residual head shifts. We determined in a few observers that, during experiments with a static TCO compensation method, where TCO is measured before and after an experiment without any ongoing monitoring, stimuli can be displaced up to 2 arcmin (Fig 7, top row). To visualize the impact of our proposed TCO compensation method, the precision of ± 0.15 arcmin determined here for our system was used to exemplary shift the target position on a retinal image due to noise of the ongoing compensation. For an experiment targeting individual cones of the central retina or rods, a maximal positional error of a single pixel, or 6 arcsec, is tolerable - a requirement that is met by about 75% of the TCO compensated positions. At 0.4 degree eccentricity, positional errors up to 12 arcsec would be acceptable. This criterion was met for all (100%) estimated TCO values in our study. Additionally, variability across three consecutively recorded calibrations in 14 tested participants fell well within 0.1 arcmin, which was lower than the observed single frame measurement noise (Fig. 4B and 5).

**Fig. 7.**
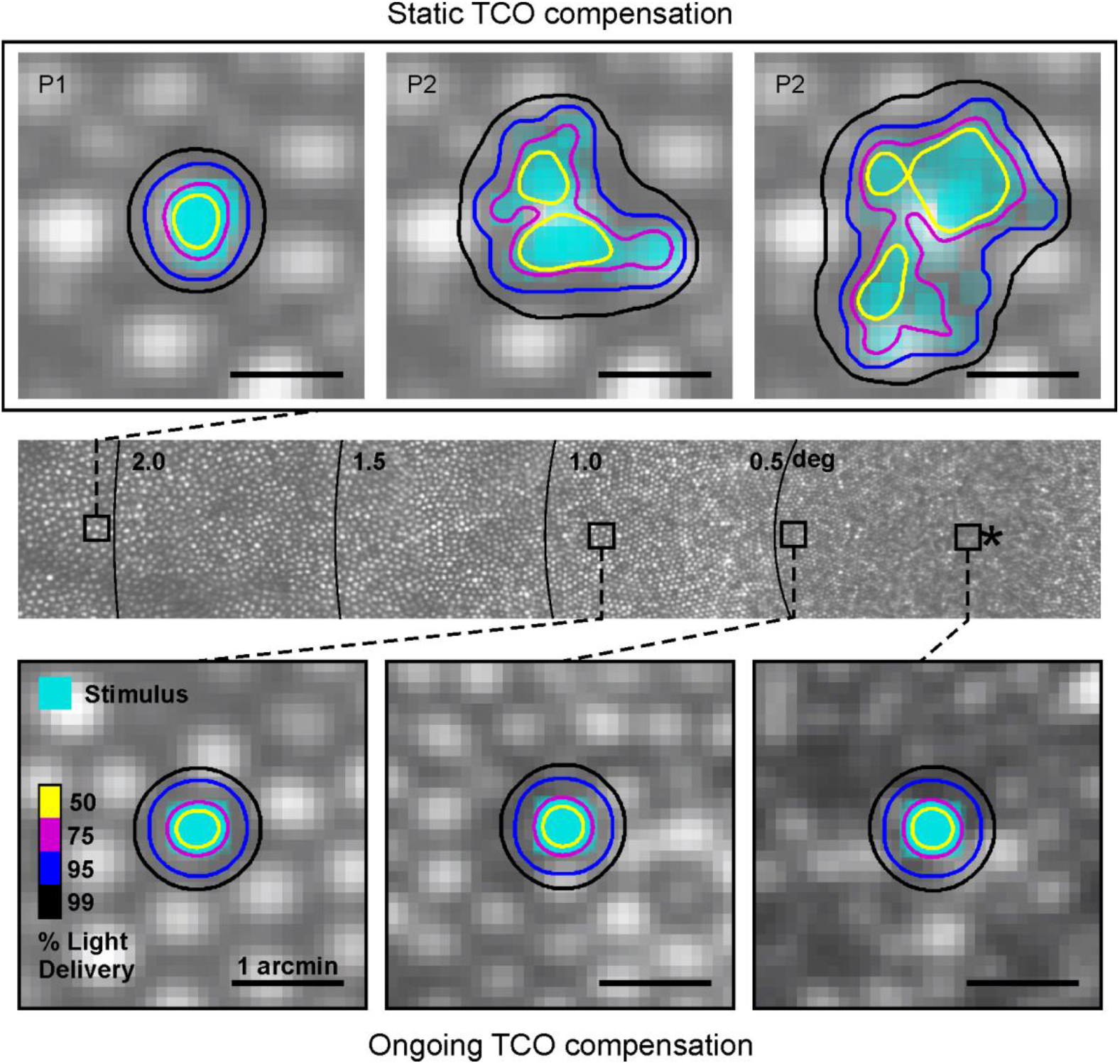
Demonstration of TCO compensation error at different retinal eccentricities. Top: Enlarged view of a typical stimulation site in AOSLO based experiments with current static TCO compensation. Overlay shows actual light delivery based on minimal head shifts in front of the system recorded by our eye tracking system during experiments in three participants (P1-3). Middle: AOSLO image montage encompassing the central fovea, asterisk marks the preferred retinal location of fixation, rings denote eccentricity. Cone size increases rapidly with increasing eccentricity. Bottom: Enlarged view of the smallest cones in the central fovea, 0.4 degree and 0.9 degree eccentricity. Overlays display theoretical stimulus delivery with ongoing TCO compensation as presented in this study. Our typical cell sized square-stimulus expands 3 pixels (diffraction limited FWHM, 0.3 arcmin).

In the subjective validation experiment, participants made an average error of 0.30 arcmin, corresponding to twice the objectively determined precision, and smaller than the expected Nyquist limit of 0.43 arcmin for a foveal cone density of 200,000 cells/mm2 [36], but slightly higher than the typically found average Vernier acuity of 0.19 arcmin [37] and 0.25 arcmin [38]. Because we did not observe any systematic displacements across the three participants, we conclude that the subjective validation confirmed the current calibration process. However, comparable to other eye tracker setups, we plan to include a brief subjective validation routine of the calibration before running single cell stimulation experiments to detect systematic mismatches between subjective and estimated TCO. These small chromatic offsets could occur, for example, due to alignment changes in the AOSLO setup.

As a practical note, for compensation of TCO during microstimulation experiments, we currently implemented a per-trial approach. Because our eye tracking software is a standalone solution, its data has to be transmitted to our stimulation software, which is run in a Matlab environment. Purkinje image coordinates were sent from the eye tracker software to Matlab on request, triggered by the participant’s keystroke that also triggers stimulation presentation. The delay of this communication was measured by means of timestamps, and did not exceed 30 msec. Based on the calculations shown above, we disregard the amount of gaze shifts for this period of time. The average head shift within a 30 msec window across all our calibration sequences was 0.006 mm, resulting in a TCO glitch of 0.02 arcmin due to transmission delays introduced by the software interface.

In summary, the demonstrated eye tracking method seems a viable solution to estimate and compensate chromatic offset in real time within a fraction of arcmin, necessary for adaptive optics microstimulation of the smallest photoreceptors of the retina. Furthermore, we experimentally confirmed the fundamental relationship between angle Kappa and TCA, which could be a convenient implementation in applications where TCA is wished to be compensated but cannot be directly measured.

